# Viability of engineered AAVs via protein language models

**DOI:** 10.64898/2026.06.11.731521

**Authors:** Mélissa Desrosiers, Tommaso Ocari, Jeanne Trinquier, Emília A. Zin, Timothé Van Meter, Maëlle Delmas, Müge Tekinsoy, Arthur Planul, Takahiro Nemoto, Deniz Dalkara, Ulisse Ferrari

## Abstract

Capsid engineering has greatly improved the performance of recombinant AAV vectors used for gene therapy. One commonly used strategy is the insertion of a short, 7-mer, peptide into surface-exposed loops to modify receptor interactions and enhance cell entry. While effective in receptor retargeting and improved transduction, these insertions might destabilize the capsid protein, hinder assembly, and thus limit production. While previous attempts have used deep mutational scanning and AI to predict which insertions are viable, there is lack in understanding the structural consequences of these peptide insertions at the amino-acid level. Here we combined experiments, deep sequencing and large protein language models to gain insight on the impact of 7-mer insertions on the VR-VIII region. We first characterize the biochemical properties of viable insertions, thus identifying which residues are well tolerated, and which should instead be avoided. We then focus on the nearby context of those insertions, by studying the effect of the linkers, either for highly diverse libraries or for individual variants known for their efficiency. Next, we study the broader context, by extending our analysis to the whole capsid sequence, and identifying regions that can tolerate insertions without long-ranged structural deformations that could affect capsid functionality. We conclude with a cross-serotype comparison and a viability analysis of tens of previously engineered variants. Our work showcases how AI can uncover structure–function rules governing the success of engineered AAV capsids.

## 1 Introduction

Adeno-associated viruses (AAVs) are among the most widely used viral vectors for in vivo gene delivery. They are currently employed in hundreds of clinical trials and have led to multiple approved gene therapies [1, 2]. Naturally occurring AAV serotypes exhibit substantial sequence and structural diversity, which translates into distinct tissue tropisms and biological properties [3]. However, these native characteristics are not necessarily aligned with specific therapeutic needs. As a result, capsid engineering has become a central strategy for expanding the functional repertoire of AAV vectors [4].

One of the most widely used capsid engineering approaches consists of inserting a short peptide, typically a seven–amino-acid sequence (7-mer), into a surface-exposed loop of the capsid, most commonly within the VR-VIII region [5–11]. This strategy enables exploration of large sequence spaces and selection of variants with altered tropism or function. However, peptide insertion is a disruptive perturbation: only a fraction of inserted sequences preserve correct folding, capsid assembly, and structural stability. Throughout this work, we refer to such variants as viable, meaning that the modified capsid proteins fold and assemble, and remain sufficiently stable to be recovered during viral production.

Understanding the determinants of insertion viability is therefore essential for interpreting directed evolution screens and for designing libraries that balance functional diversity with productive yield [12]. Several experimental and computational efforts have begun to address this challenge, including comprehensive mutational scans [13] and supervised machine-learning models trained on serotype-specific insertion datasets [12, 14–16]. These previous studies have primarily mapped substitution or insertion tolerance in a serotype-specific manner and provided predictive tools for particular datasets. However, they largely treat insertion sites as isolated classification problems and offer limited insight into how inserted peptides perturb the biochemical and structural context of the full capsid protein.

Here we combine new results from in house viral library productions, previously published deep sequencing datasets, and large protein language models (PLMs) [17] to study how the amino-acid content of inserted 7-mers impacts the overall capsid protein. Unlike previous predictive approaches that rely primarily on labeled insertion datasets, PLMs learn representations from evolutionary sequence data that encode long-range biochemical and structural dependencies across entire proteins. This allows us to analyze insertions in their full capsid context rather than as site-specific classification tasks, making our approach qualitatively different from prior models. Rather than focusing exclusively on prediction accuracy, we use these models to extract interpretable biochemical and structural constraints governing 7-mer insertion viability.

We show that 7-mer insertion viability at the VR-VIII insertion site is largely determined by simple amino-acid composition rules, with additional contributions from linker effects and local structural context. By extending these analyses capsid-wide, we identify regions that combine high insertion tolerance with strong structural locality. Importantly, this goes beyond the conventional definition of capsid variable regions, which are typically identified based on sequence diversity or evolutionary variability. Instead, our approach resolves quantitative, position- and serotype-specific constraints that determine whether an inserted peptide preserves folding, assembly, and capsid stability. Finally, we show that insertion tolerance at the VR-VIII insertion site is transferable across serotypes: when local backbone geometry is conserved and insertion effects remain local, a model calibrated on one serotype preserves substantial predictive ranking power on others. Together, these results establish a practical framework for interpreting and designing peptide insertions in AAV capsids that is applicable even when experimental insertion data remain limited.

## 2 Results

### Amino acid composition dominates 7-mer insertion viability

To understand how short peptide insertions affect AAV capsid assembly, we analyzed a collection of eight 7-mer insertion libraries generated across multiple laboratories, including published AAV9 datasets [9–11, 16], together with new AAV2 and AAV5 libraries produced in this work. (Methods Section 4.1 and Tables 1 and 2.)

**Table 1:** Per-variant datasets used for supervised model training and evaluation. *Variants* is the number of unique insertion variants, *HEK cells* is the total producer-cell count, and *Cells/variant* is the average cellular coverage per variant.

| Serotype | Use | Variants | HEK cells | Cells/variant | Reference |
| --- | --- | --- | --- | --- | --- |
| AAV2 | Training / Testing | 53,382 | 100,000,000 | 1,873.29 | In-house dataset |
| AAV5 | Training / Testing | 737,587 | 100,000,000 | 135.58 | In-house dataset |
| AAV9 | Training / Testing | 68,776 | 660,000,000 | 9,596.37 | Eid et al. [16] |

**Table 2:** Broad-coverage datasets used for amino-acid enrichment analyses and aggregate comparisons. The in-house AAV2 viability-selection datasets from rounds 1 and 2 are used in Fig. 5b.

| Serotype | Use | Reference |
| --- | --- | --- |
| AAV2 | AA enrichment (Supplementary Fig. S1); viability selection rounds 1 and 2 (Fig. 5b) | In-house dataset |
| AAV2 | AA enrichment (Supplementary Fig. S1) | Byrne et al. [41] |
| AAV9 | AA enrichment (Supplementary Fig. S1) | Stanton et al. [11] |
| AAV9 | AA enrichment (Supplementary Fig. S1) | Tabebordbar et al. [10] |
| AAV9 | AA enrichment (Supplementary Fig. S1) | Ravindra Kumar et al. [9] |

All libraries follow a common experimental strategy in which a foreign 7-mer peptide is inserted into a solvent-exposed capsid loop, and the resulting variant populations are quantified by deep sequencing to assess capsid assembly and production. In all cases, the insertion is made within the VR-VIII surface loop at the capsid’s threefold symmetry axis (insertion site at residue 588 in AAV2, residue 576 in AAV5, and residue 589 in AAV9), which forms part of the receptor-binding region involved in cell entry. Because of both its functional relevance and accessibility on the capsid surface, this site has become the standard locus for peptide display and retargeting [5–11].

Empirical viability was defined as the log ratio between a variant’s abundance in the viral pool and its abundance in the plasmid library (Fig. 1a; Methods, Section 4.3). Viability distributions were sharply bimodal (Fig. 1b), with most insertions being either clearly tolerated or strongly disruptive. This switch-like behavior indicates that the loop can accommodate certain biochemical changes but is highly sensitive to others.

**Figure 1:**
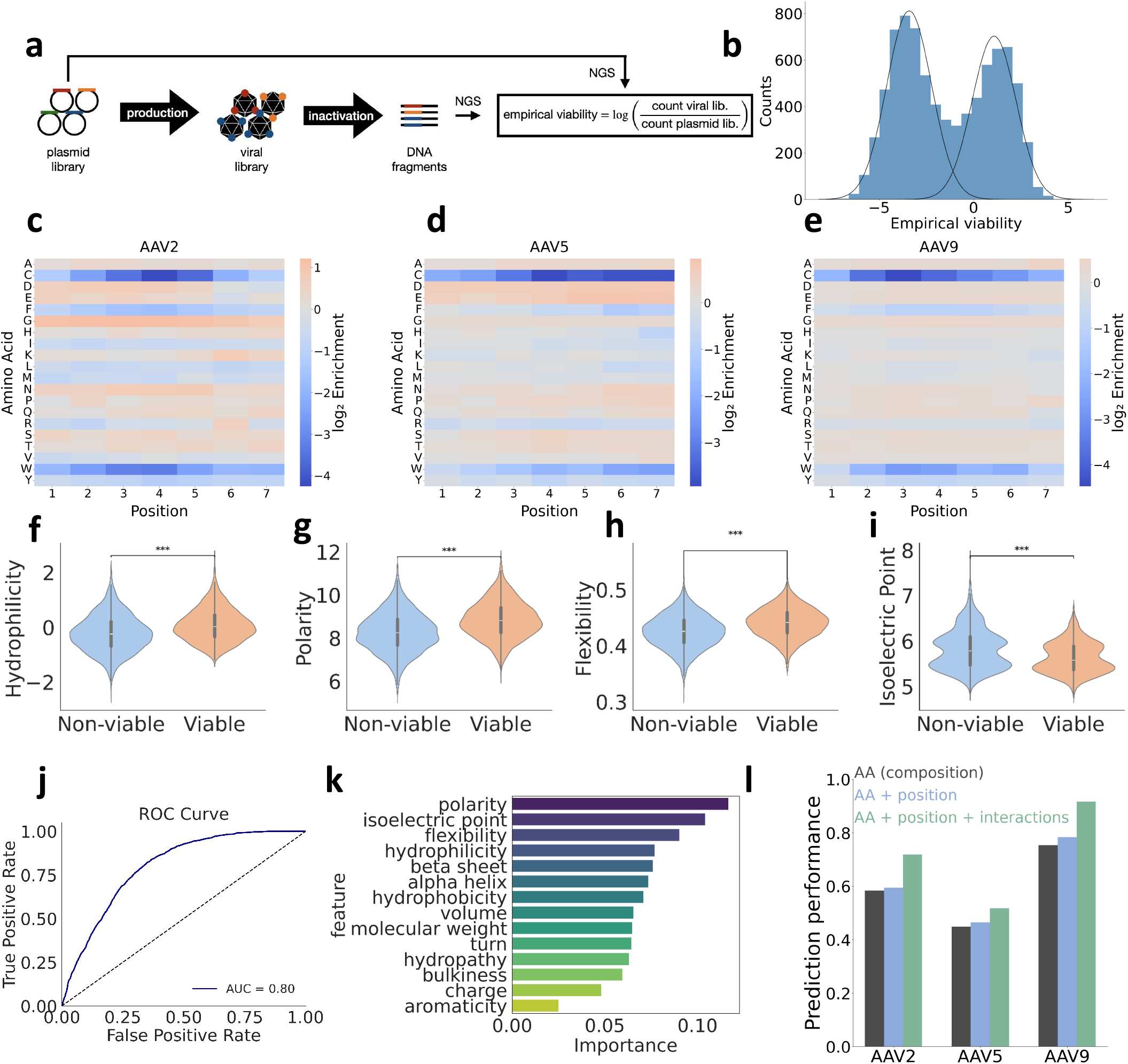
Amino acid composition dominates the determinants of AAV insertion viability. **a**, Schematic of the experimental workflow. A plasmid library encoding randomized 7-mer peptide insertions was packaged into AAV capsids to generate a corresponding viral library. Both plasmid and viral pools were sequenced by next-generation sequencing (NGS). Empirical viability was computed as the log ratio of viral to plasmid read counts, corrected for viral inactivation. **b**, Bimodal distribution of empirical viability values for the AAV2 dataset. The left peak corresponds to non-viable variants, while the right peak corresponds to viable insertions. **c to e**, Empirical log-enrichments of the 20 amino acids at each of the seven inserted positions for AAV2, AAV5, and AAV9 libraries, respectively. **f to i**, Violin plots comparing biophysical properties, hydrophilicity (**f**), polarity (**g**), flexibility (**h**), and isoelectric point (**i**), between viable and non-viable variants. **j**, Classification performance (AUC) of a random forest model trained on these biophysical features. **k**, Feature importance scores from the random forest, highlighting the relative contribution of each property. **l**, Pearson correlation between predicted and empirical viability for different model architectures, ranging from amino acid composition only to models incorporating positional and interaction information within the 7-mer insert.

Amino-acid enrichment analyses revealed highly reproducible insertion preferences across AAV2, AAV5, and AAV9 (Fig. 1c–e), as well as across additional datasets generated by independent laboratories (Supplementary Fig. S1a–e). Bulky or hydrophobic residues (F, W, Y) and cysteine (C) are consistently depleted among viable variants, whereas small, flexible, or polar residues are enriched. In the enrichment heatmaps (Fig. 1c–e), these depletion/enrichment patterns are broadly similar across the seven 7-mer positions (i.e., weak column-to-column variation), indicating that overall biochemical composition, rather than precise residue order, is the dominant determinant of viability.

Consistent with this observation, viable insertions tend to be more flexible, hydrophilic, and polar, whereas non-viable insertions tend to be more rigid or hydrophobic (Fig. 1f–i; Methods, Section 4.5). In these violin plots, each point corresponds to one 7-mer, after averaging the property across its seven residues. The distributions of viable and non-viable variants overlap substantially, but they are shifted in a consistent direction across properties. Statistical significance was assessed with a two-sided Mann–Whitney U test, a non-parametric test that compares whether two independent groups tend to have different values.

To test whether these coarse biochemical properties are informative on their own, we trained a random-forest classifier on the averaged properties (Methods, Sections 4.4 and 4.7). A random forest combines many simple decision rules into a single classifier and asks whether viable and non-viable insertions can be separated from these features alone. Performance was measured by the area under the receiver operating characteristic curve (AUC), which summarizes how well the two classes can be distinguished across all decision thresholds.

Using this approach, the classifier separates viable from non-viable variants well across all serotypes (Fig. 1j). Feature-importance analysis (Fig. 1k) highlights hydrophilicity, polarity, and flexibility, although these properties are themselves correlated and should not be interpreted as fully independent determinants. Together, these results reinforce the idea that viability is driven largely by broad biochemical tendencies rather than by detailed sequence motifs.

Finally, to directly assess the contribution of sequence order and residue–residue interactions beyond simple composition, we trained a hierarchy of sequence-based models of increasing complexity to predict empirical viability directly from 7-mer sequences (Methods, Section 4.7). A position-independent model based solely on amino-acid composition already performs well, while allowing position-specific effects or nonlinear interactions yields only small gains in predictive performance (Fig. 1l). Differences in absolute predictive performance across serotypes (for example, lower correlations for AAV5 than for AAV9) primarily reflect differences in experimental noise and library quality, consistent with the lower Cells/variant values in Table 1, rather than intrinsic biological differences.

Taken together, these results define the biochemical constraints governing 7-mer insertions at the VR-VIII surface loop and demonstrate that amino-acid composition, rather than precise sequence patterning, is the primary determinant of insertion viability across AAV serotypes and experimental platforms.

### Rationalizing linker design using a protein language model

Insertions at the VR-VIII surface loop are typically flanked by short *linker* sequences whose dual purpose is to preserve local structural compatibility while maintaining adequate exposure of the inserted peptide on the capsid surface. Despite this functional importance, linker design in AAV engineering has remained largely heuristic. Laboratories typically rely on simple motifs such as `LA-A` [7, 18–20] or `AAA-AA` [5, 21–23], motivated by their presumed flexibility and minimal steric impact, yet without a quantitative framework to justify or refine these choices. As a result, linker selection is rarely optimized or even interrogated, and its contribution to insertion viability is often implicitly assumed rather than explicitly evaluated. A comprehensive, linker-focused mutagenesis scan would be highly combinatorial across inserts and linker architectures; here we instead used zero-shot PoET [17] to perform a controlled *in silico* scan that prioritizes linker motifs and generates testable hypotheses for future validation (Methods, Section 4.8). PoET has been shown to perform exceptionally well on AAV capsid mutational data, ranking among the top models in the ProteinGym benchmark [24] and showing strong agreement with our experimental measurements (Supplementary Table S1). This makes it a suitable tool for extracting sequence-level constraints in regions such as linker segments that lack dedicated experimental mutagenesis datasets.

At the library level, we asked which linker residues would be most broadly favorable when building a starting directed-evolution library that contains many different inserts. To address this, we generated simulated libraries in which each of 500 distinct 7-mer insertions was paired with 1,000 alternative linker sequences. We then used PoET to score all of these constructs and averaged the results across inserts, so that the final enrichment profiles capture linker preferences that are broadly compatible with many sequence contexts rather than optimized for a single variant. The resulting heatmaps for `XX-X`, `XXX-XX`, and `XX-XXX` linkers (Fig. 2a–c) reveal consistent biochemical rules: residues that are small, flexible, and polar are broadly favored, whereas bulky hydrophobic or aromatic residues are generally disfavored. The same library-level analyses performed separately for AAV5 and AAV9 showed the same qualitative trends (Supplementary Fig. S2). More specifically, PoET predicts strong enrichment for **A, G, P, T, S, N, and Q** across most linker positions. In contrast, **C, W, F, Y, and M** are strongly depleted, consistent with their steric bulk and reduced conformational flexibility. Charged residues show distinct behavior: acidic residues (**D, E**) are mildly tolerated, whereas basic residues (**K, R**) tend to be disfavored. These residue-level rules are also visible when entire linker motifs are compared (Fig. 2d–f). Some linkers that are commonly used in the literature already perform well in our simulations. For example, `AAA-AA` remains among the best motifs in longer-linker settings, and the published motif `TG-GLS` [12] also falls in a high-scoring part of the landscape. By contrast, `LA-A` is robust but not always optimal: in shorter-linker settings, alternatives such as `TA-T`, `TT-D`, and `TG-S` receive higher predicted scores. Taken together, these results suggest that commonly used linkers often capture the right biochemical logic, but that better choices may still exist. We next asked whether this same logic can help evaluate the linker choice for an already engineered variant (Fig. 2h–i). To do this, we curated 25 previously reported AAV2 insertion variants (Supplementary Table S2) [5, 7, 18–23] and compared the experimentally used linker against 10,000 in silico alternatives of the same architecture. In most cases, the literature linker already ranks near the top of the predicted distribution. This is not surprising, since these variants were reported precisely because they worked experimentally, and their linker choice was therefore at least compatible with a successful outcome. However, a small subset of inserts, most notably `KDPKTTN`, `HQDTTKN`, and `ISDQTKH` [7, 20], rank much more poorly with `LA-A`. These sequences are rich in polar and charged residues, suggesting that some inserts are much more sensitive than others to linker composition. In such cases, PoET identifies alternative linkers with substantially better predicted compatibility, providing concrete, sequence-specific opportunities for optimization.

**Figure 2:**
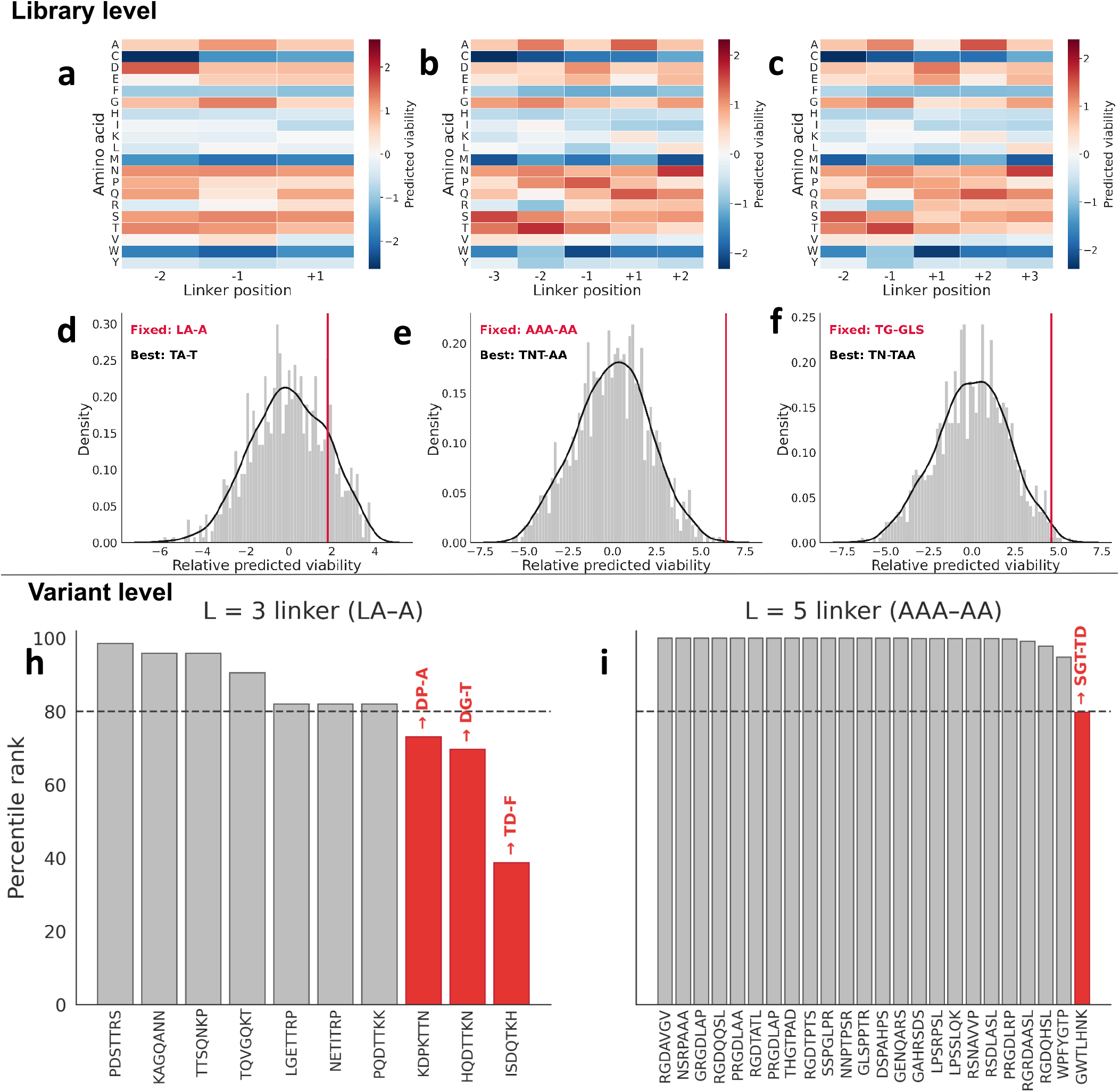
PoET-guided rationalization of linker choice at the library and variant levels. **a–c**, PoET-predicted amino-acid log-enrichment heatmaps for linker layouts `XX-X`, `XXX-XX`, and `XX-XXX`. Each heatmap reports position-specific residue enrichments aggregated over simulated libraries containing 500 distinct 7-mer inserts, each paired with 1,000 linker candidates. **d–f**, Histograms of mean predicted viability for the corresponding linker libraries, summarizing linker-level score distributions for each layout. Vertical lines indicate literature-used motifs, including `LA-A` [7, 18–20], `AAA-AA` [5, 21–23], and `TG-GLS` [12]. **h–i**, Variant-level evaluation of linker performance for literature-derived 7-mer insertions. For each curated insertion, the percentile rank of the experimentally used linker is computed against 10,000 in silico alternatives of matched linker architecture. Insertions for which the literature linker falls below the 80th percentile are highlighted as potentially suboptimal (an arbitrary threshold chosen for visualization).

Together, this analysis demonstrates how a protein language model can transform linker design from an adhoc heuristic into a quantitatively grounded step. In practice, PoET can guide linker choice at two scales: at the library level, it can help design linkers that do not depress baseline viability across diverse inserts; and at the variant level, it can suggest insert-specific linkers that preserve capsid performance and gene-delivery efficiency.

### Capsid-wide analysis of insertion tolerance and structural locality

Having established the biochemical constraints that govern 7-mer insertion viability and the local sequence preferences of the linker-loop environment, we next asked how these observations extend to the scale of the full capsid (Fig. 3). This is relevant not only for identifying sites that may support future engineering, but also because different exposed regions of the capsid contribute to distinct aspects of AAV biology, including receptor binding, intracellular trafficking, and capsid stability [25–29]. Identifying insertion sites beyond the best studied loop may therefore open access to functional space that cannot be reached by repeatedly engineering VR-VIII alone. We focused here on 7-mer insertions because short peptides of this size are a standard format in AAV capsid engineering, and they represent a substantially stronger perturbation than a single amino-acid insertion while remaining directly relevant to library design. Because a truly capsid-wide experimental map of 7-mer insertions would require dense coverage of hundreds of insertion positions and extensive sampling of the 20^7^ sequence space (with sufficient depth and replication to obtain stable position-resolved estimates), we use PoET as an *in silico* probe of positional constraints. As a sanity check before applying PoET to hypothetical 7-mer insertions, we first asked whether it recovers the broad positional trends seen in the closest available experimental benchmark, namely single amino-acid insertions measured across the AAV2 capsid.

**Figure 3:**
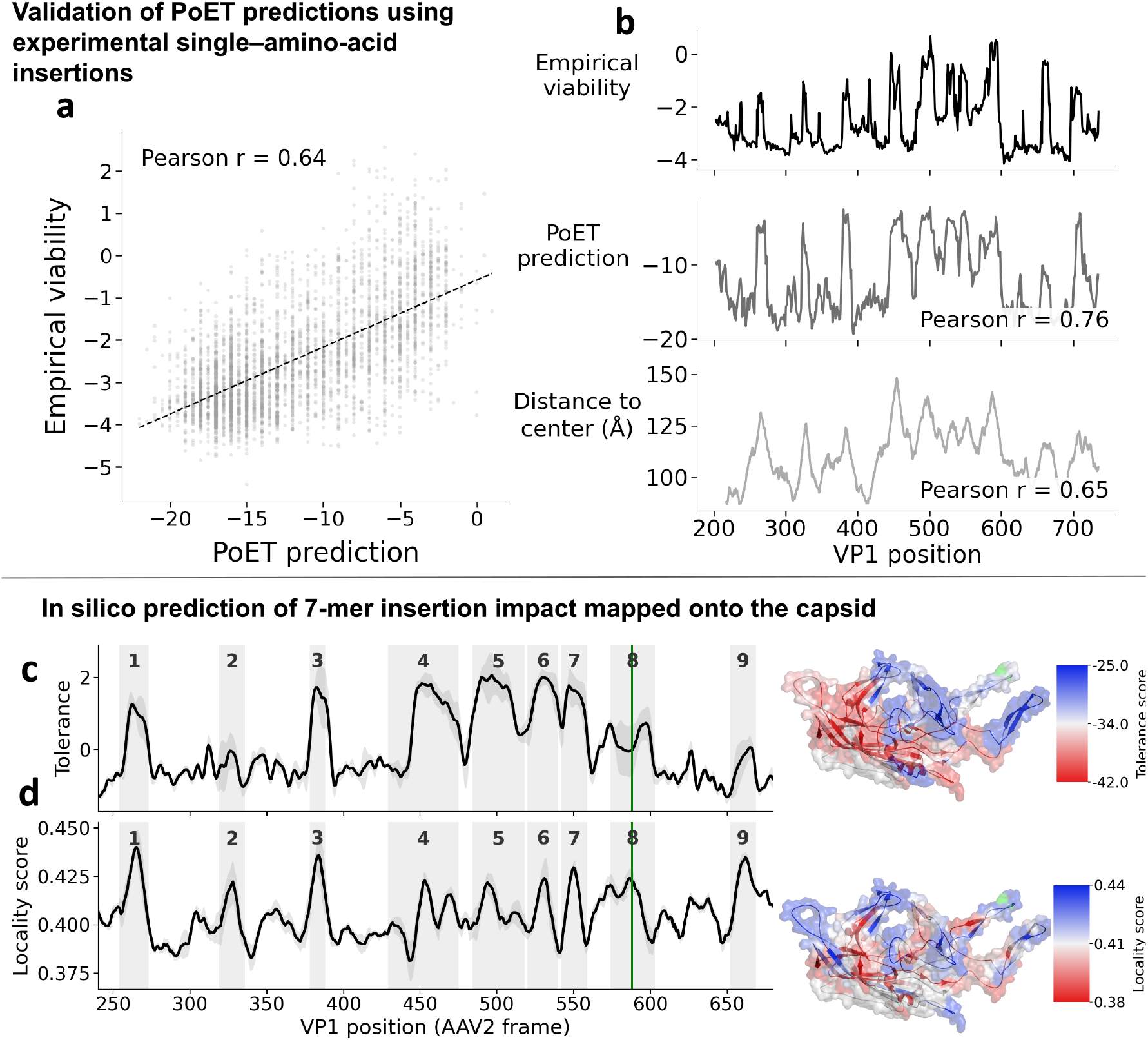
Validation and capsid-wide application of PoET for insertion tolerance prediction. **a**, Scatter plot comparing PoET predictions for single–amino-acid insertions with empirical insertion viability measured experimentally [13]. Each point corresponds to one variant, and the Pearson correlation coefficient reports the global agreement between PoET predictions and experimental measurements. **b**, Mean empirical insertion viability per VP1 position (top), compared with the corresponding mean PoET prediction (middle) and a geometric baseline given by the distance to the capsid center (bottom). Pearson correlation coefficients quantify agreement with the empirical profile for PoET predictions and for the distance-based baseline. **c**, Capsid-wide prediction of 7-mer insertion tolerance using PoET, computed from the mean change in zero-shot PoET log-likelihood upon inserting random 7-mers at each position (shown as a *z*-score across VP1 positions), and plotted as the mean profile across serotypes with standard-deviation bands. Variable regions are shaded in gray, and the experimentally targeted insertion position is highlighted in green. The capsid structure is colored by tolerance score, with low-tolerance regions shown in red and highly permissive regions shown in blue. **d**, Locality score along VP1, defined as the fraction of the predicted structural response confined within a local ±5-residue window around the insertion site, shown as the mean profile across serotypes with standard-deviation bands. Variable regions are shaded in gray, and the experimentally targeted insertion position is highlighted in green. Higher locality values indicate perturbations that remain spatially localized rather than propagating through the capsid. The capsid structure is colored by locality score using the same colormap, highlighting regions that are structurally insulated from global deformation. Green shaded regions correspond to positions jointly selected for high insertion tolerance and high structural locality, identifying capsid sites that are both permissive to diverse insertions and predicted to minimize long-range structural perturbations.

We began by validating zero-shot PoET predictions against available experimental data for single amino-acid insertions. Figure 3a compares PoET-predicted insertion compatibility with empirical viability measurements reported previously [13]. PoET predictions show good global agreement with experimental outcomes (Pearson *r* = 0.64), indicating that the model captures positional constraints that are relevant to experimentally observed insertion viability. This comparison provides a useful check that PoET recovers the same broad capsid regions that are more or less permissive to insertion experimentally.

To contextualize this agreement along the capsid sequence, we next examined how insertion tolerance varies across VP1. Figure 3b shows the mean empirical insertion viability per position alongside two predictive baselines: the corresponding mean PoET prediction and a purely geometric proxy given by the radial distance of each residue from the capsid center. While radial distance correlates with empirical viability (Pearson *r* = 0.65), PoET predictions show substantially higher agreement (Pearson *r* = 0.76), indicating that PoET captures structural determinants beyond simple geometry.

We next used PoET to explore a regime that remains largely inaccessible experimentally: a capsid-wide *in silico* survey of hypothetical 7-mer insertions across VP1 (Fig. 3c,d). For each position, we simulated 100 random 7-mer insertions and extracted two complementary quantities characterizing the capsid’s response (Methods, Section 4.8). The first quantity, the **insertion tolerance score** (Fig. 3c), quantifies the mean change in zero-shot PoET log-likelihood induced by random 7-mer insertions at that position (reported as a *z*-score across VP1 positions for visualization). This analysis reveals a highly heterogeneous tolerance landscape, with buried or geometrically constrained regions strongly disfavoring insertions, and several surface-exposed loops exhibiting moderate to high tolerance.

Importantly, tolerance alone does not distinguish whether an insertion perturbs only the local loop or induces broader structural rearrangements. The insertion of a 7-mer peptide can affect capsid function not only by destabilizing folding or assembly, but also by triggering long-range deformations that alter receptor engagement, endosomal escape, uncoating, or nuclear import. To capture this distinction, we computed a second metric, the **structural locality score** (Fig. 3d), defined as the fraction of the predicted structural response confined within a local ±5-residue window around the insertion site. Jointly considering tolerance and locality identifies a limited subset of positions that combine permissiveness with structural insulation (green shaded regions in Fig. 3c,d).

To evaluate how these trends vary across capsid backgrounds, we computed the same tolerance and locality profiles for additional serotypes (Supplementary Fig. S3). While the broad biochemical logic is conserved, the absolute profiles are serotype specific, indicating that insertion-permissive windows are not identical across AAV backgrounds. This serotype dependence may help explain why peptide-display strategies do not always transfer equally well between capsids, including weaker transfer to backgrounds such as AAV8 in pan-AAV settings [30].

Within this landscape, the VR-VIII insertion site emerges as a favorable compromise between these two properties. Some positions score higher in tolerance alone, but fewer combine relatively high permissiveness with a similarly local predicted response. We therefore view VR-VIII as a useful reference point rather than a unique exception: it illustrates the type of balance that may help prioritize candidate grafting sites across the capsid.

### Cross-serotype generalizability of insertion tolerance at the VR-VIII insertion site

Many AAV serotypes have been identified in nature [3], and new variants continue to be discovered [31, 32]. Yet even when different serotypes are engineered at structurally analogous sites in the threefold-axis loop [33], they do not tolerate inserted peptides equally well: the same 7-mer can be highly compatible in one capsid background and much less so in another. This raises a practical question for capsid engineering: even if absolute compatibility shifts across serotypes, is relative ranking conserved well enough that data from one serotype can help prioritize inserts in another? In practice, such ranking information is often the most useful quantity for experimental design, because it enables library construction and screening strategies that enrich for highly viable candidates while still preserving diversity for downstream functional selection.

Because insertion effects at VR-VIII are largely local (Fig. 3) and the loop geometry is similar among AAV2, AAV5, and AAV9 (Fig. 4b) [33], we hypothesized that transferable insertion rules should exist at this site even across phylogenetically distant serotypes (Fig. 4a). To test this, we used PoET embeddings as a shared representation and trained a lightweight supervised head on top of the frozen embeddings from the seven inserted residues at the serotype-specific VR-VIII site (residue 588 in AAV2, 576 in AAV5, and 589 in AAV9). This supervised calibration improves agreement with empirical viability compared to zero-shot scoring (Supplementary Table S1; Methods, Section 4.9), and we refer to the resulting model as AAV-InsertPoET.

**Figure 4:**
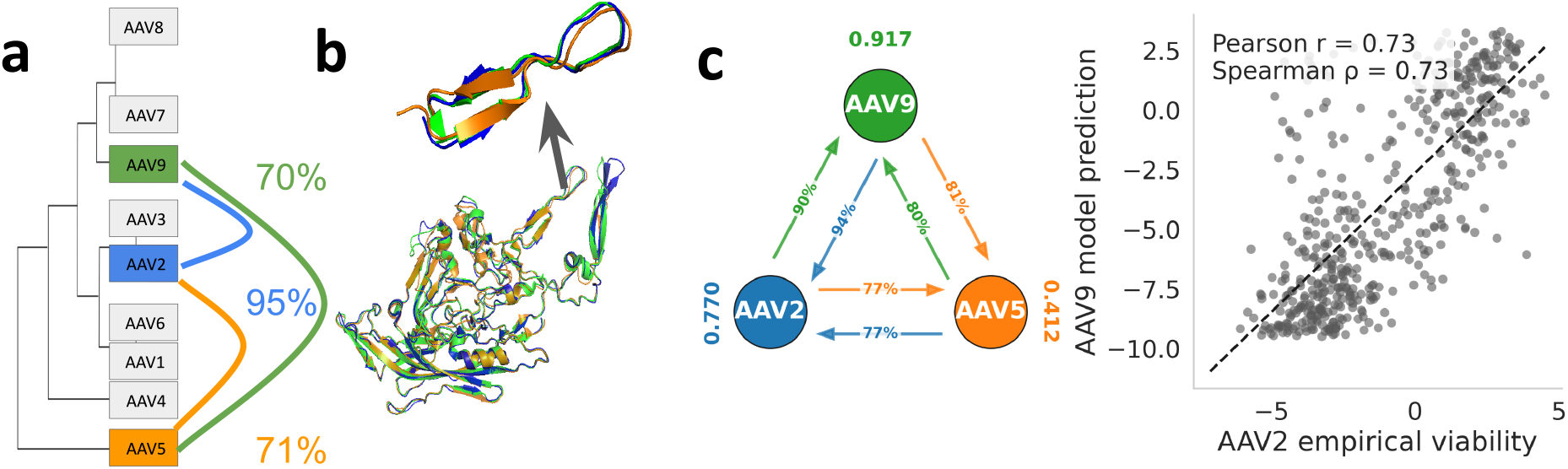
Cross-serotype generalization of insertion tolerance models at the VR-VIII insertion site. **a**, Phylogenetic relationships between AAV serotypes based on VP1 structure similarity, with percent identities shown on the right. **b**, Structural alignment of VP1 monomers from AAV2, AAV5, and AAV9, highlighting the aligned VR-VIII insertion loop (AAV2 residue 588-equivalent; insertion sites at residue 588 in AAV2, 576 in AAV5, and 589 in AAV9). **c**, Cross-serotype prediction of empirical insertion viability with AAV-InsertPoET. Arrows summarize train-to-test transfer performance in percent, and the scatter plot shows AAV9 to AAV2 transfer.

Within each serotype, AAV-InsertPoET accurately recovers empirical viability on held-out data, providing a quantitative baseline for transfer. We then evaluated cross-serotype transfer by training on one serotype and testing on another (Fig. 4c). Transfer between AAV2 and AAV9 is strongest in both directions, consistent with shared local constraints at VR-VIII. Transfers involving AAV5 are weaker but remain clearly informative, indicating that useful ranking information still carries across more divergent capsid backgrounds. Together, these results show that cross-serotype transfer can be practically useful for prioritizing candidate inserts in new capsid backgrounds: absolute tolerance remains serotype specific, but relative ranking is sufficiently conserved to support transfer.

### Analysis of literature insertions

Starting from different wild-type AAV serotypes, the insertion of short peptide motifs has enabled the engineering of numerous capsid variants with improved functional properties, including altered tropism, receptor usage, and immune evasion. Most such approaches rely on directed evolution strategies involving repeated rounds of viral production and selection [5, 7, 9, 11, 18–23, 34–40]. Because capsid assembly and viral production are prerequisites for all downstream assays, it is important to quantify how strongly viability alone shapes the outcome of these selection workflows.

To address this question, we compiled a curated set of engineered AAV variants containing 7-mer peptide insertions reported in the literature (Supplementary Table S2) and evaluated their predicted viability using AAV-InsertPoET. We compared these scores to those of both random 7-mer insertions and variants enriched after one or two rounds of viability-based selection (Fig. 5). Selected variants are taken from our in-house AAV2 production-only selection dataset (Methods, Section 4.1; Table 2).

**Figure 5:**
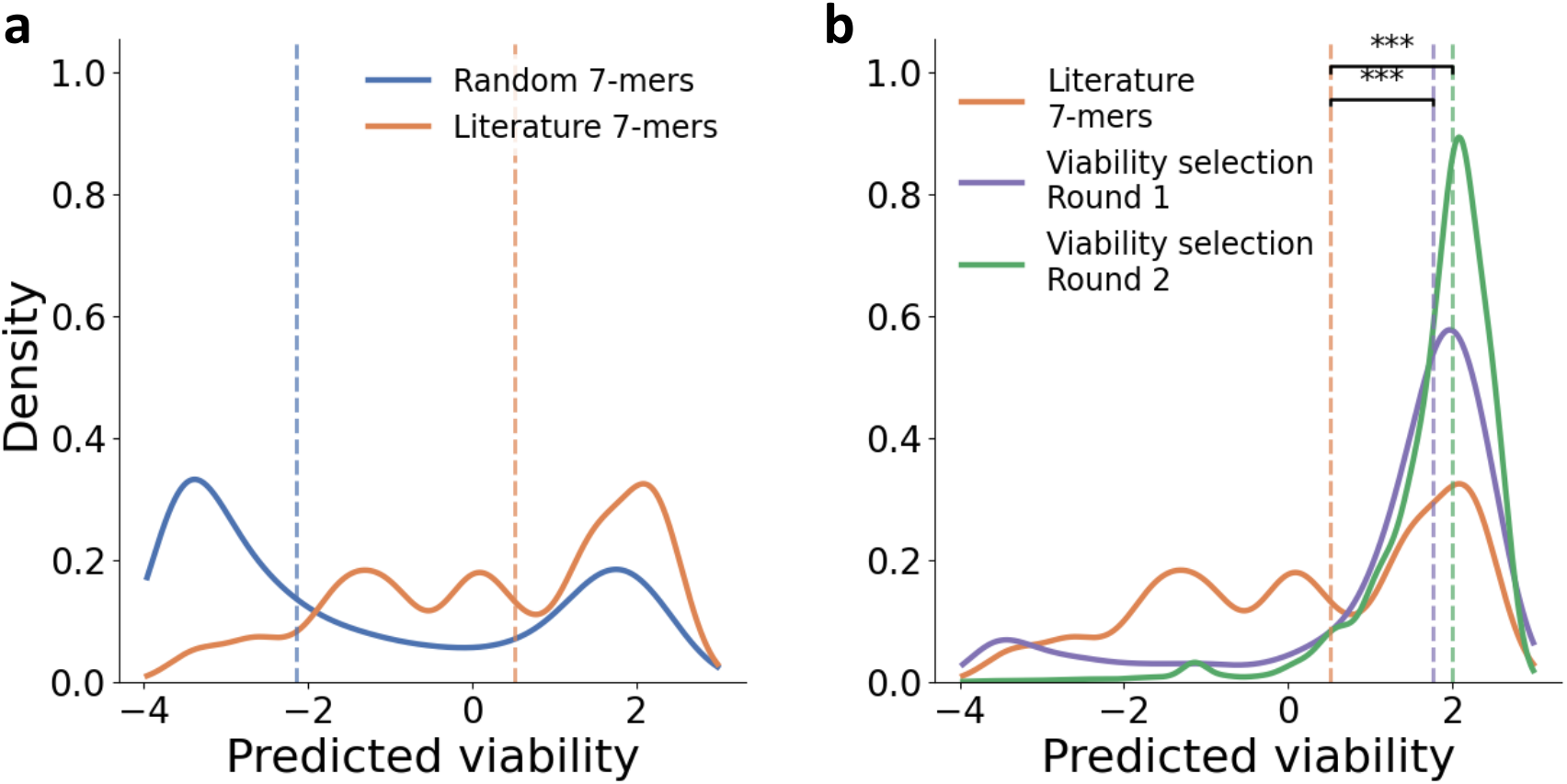
PoET viability predictions for random, selected, and literature 7-mer insertions. **(a)** Distribution of predicted viability scores for random 7-mer insertions (blue) and engineered 7-mer insertions reported in the literature (orange). **(b)** Distribution of predicted viability scores for literature 7-mers (orange) and variants enriched after one (purple) or two (green) rounds of viability selection. Vertical dashed lines indicate the median of each distribution.

Literature-reported insertions exhibit substantially higher predicted viability than random 7-mers (Fig. 5a; median 0.52 vs. −2.15).

Variants enriched solely through viral production exhibited substantially higher predicted viability than literature variants (Fig. 5b). After one round of selection, the median predicted viability increased to 1.77, compared to 0.52 for literature variants, corresponding to a KS distance of *D* = 0.33 and a bootstrap *p*-value of 8.3 × 10^−4^. After two rounds, the shift was further amplified (median 2.00; *D* = 0.46, *p* = 8. 3 ×10^−4^), indicating a strong and statistically significant global redistribution of viability scores. Together, these results show that while engineered variants reported in the literature are clearly biased toward higher viability relative to random insertions, they do not reach the extreme viability levels achieved by variants selected solely for efficient production. Therefore, we conclude that published variants are shaped by additional constraints beyond production viability, consistent with multi-objective selection.

## 3 Discussion

In AAV capsid engineering, peptide insertions that successfully improve tropism or receptor usage represent only a fraction of all variants tested, as a large proportion fail upstream due to an inability to assemble into capsids that can be efficiently produced. To overcome this challenge in capsid engineering, we provide a quantitative framework to characterize 7-mer insertion viability at the VR-VIII insertion site across AAV serotypes. Combining curated viability datasets with protein language model representations, we identify robust composition-driven rules, linker effects, and position-specific structural constraints that govern successful insertions.

We find that viability at the VR-VIII insertion site is governed primarily by the biochemical composition of the inserted 7-mer, whereas the precise residue order has a more limited effect. The fact that these compositional biases are shared across serotypes suggests that even a simple smart library could already enrich for producible variants by avoiding the most strongly disfavored amino acids [12, 16].

Inserted peptides in the VR-VIII region are typically flanked by short linker sequences, whose role is both to preserve local structural compatibility and to maintain sufficient exposure of the inserted peptide at the capsid surface. In line with this, our results show that linker composition contributes directly to viability, and that this effect is not random: using the protein language model PoET [17], we recover broad residue-level rules, identify linker motifs that remain robust across many inserts, and reveal sequence-specific exceptions. The novelty here is that protein language models provide a way to rationalize linker choice, a design parameter that is widely used in engineered AAV variants but is usually chosen empirically rather than justified explicitly from sequence or structural constraints [7, 12, 20]. For experimentalists, this offers a practical tool both for choosing an initial linker and for deciding whether linker optimization is worth revisiting after a promising inserted sequence has already been identified.

Beyond the immediate sequence context of the insertion, our results show that the same approach can be extended to the scale of the whole capsid, where it provides a more quantitative view of insertion permissiveness than the classical notion of AAV variable regions alone. Our results move beyond this descriptive view by defining two quantitative scores, one for insertion tolerance and one for structural locality. The first measures how compatible a given region is with the presence of a foreign peptide, whereas the second measures whether the predicted structural response remains confined near the insertion site or instead propagates more broadly through the capsid. This distinction is important because different exposed regions of the capsid contribute to different aspects of AAV biology, including receptor binding, intracellular trafficking, and capsid stability [25–29]. In that sense, these two scores provide a principled way to identify new insertion sites beyond VR-VIII, which is valuable not only for expanding the repertoire of engineerable capsid positions, but also for exploring biological functions that may not be accessible from the best studied insertion loop alone.

Compared with previous approaches, our analysis provides a unified way to reason about insertion permissiveness at the scale of the whole capsid. Existing capsid-wide experimental maps are, for now, restricted to a single serotype and to simpler perturbation regimes such as single amino-acid insertions [13], while insertion-prediction and library-design studies typically rely on supervised models trained on labeled datasets generated in a specific serotype and insertion context, most often at a single loop [12, 15, 16]. By contrast, the tolerance and structural-locality scores introduced here provide a quantitative, sequence-based view that can be applied across positions without requiring a dedicated mutagenesis map for each site.

When experimental viability data are available, this framework can be extended from unsupervised analysis to supervised prediction. In our case, calibrating PoET on AAV2, AAV5, and AAV9 provides accurate serotype-specific models and, importantly, shows that part of insertion viability is shared across serotypes, since relative ranking remains partly conserved even when absolute compatibility shifts. This is particularly useful for experimentalists, because it means that data from a well-characterized capsid can help prioritize variants in another background where viability remains poorly mapped. More broadly, it also illustrates a practical advantage of protein language model representations: the same learned sequence representation can be reused across related insertion problems, rather than building a separate predictor from scratch for each serotype or loop. In that sense, our approach complements previous supervised studies, which were trained in a specific serotype and insertion context [12, 15, 16].

Our analysis of published insertion variants further shows that viability is an important but incomplete determinant of engineering success. Published variants are clearly enriched for viability relative to random 7-mers, but they remain less extreme than variants recovered under production-only selection. This suggests that the successful variants reported in the directed-evolution literature are shaped by at least two constraints: they must remain viable, but they must also satisfy the downstream property under selection [5–7, 9–11, 39]. In many cases, however, these contributions are not disentangled, so it remains unclear whether the final variants are truly optimized for the target task or instead reflect a strong bias toward viability. This may help explain why some directed-evolution strategies fail or yield only modest gains, especially when the starting library or the selection scheme imposes viability constraints that are too strong. In this context, our results are encouraging because they show that viability can already be modeled with good predictive accuracy, and that part of this signal generalizes across serotypes. This provides a practical basis for designing starting libraries that preserve functional diversity while removing clearly non-viable variants, and more broadly for explicitly separating viability from downstream selectivity in future modeling efforts.

## Acknowledgments

The authors would like to thank L. C. Byrne for providing the AAV dataset 2, J. Fernandez-de-Cossio-Diaz, G. Uguzzoni, and L. C. Byrne for useful comments and discussions. The authors thank Twist Bioscience for producing the plasmid library and the platform GENOMIC at Institut Cochin and at Institut Pasteur for NGS sequencing. This work was supported by DIM C-BRAINS, funded by the Conseil Régional d’Íle-de-France; ERC Starting Grant (REGE-NETHER 639888 to D.D.); European Research Council (ERC) Horizon 2020 Framework Programme Project (863214 – NEUROPA to D.D.); UN-ADEV; BpiFrance (Grant i-Demo – GEAR project to D.D. and U.F.); the Institut National de la Santé et de la Recherche Médicale (INSERM); Sorbonne Université (to D.D. and U.F.); The Foundation Fighting Blindness; Agence Nationale de la Recherche (ANR) RHU Light4-Deaf; LabEx LIFESENSES (ANR-10-LABX-65 to D.D.); IHU FOReSIGHT (ANR-18-IAHU-01 to D.D.); Paris Region Postdoctoral Fellowship (PRPF to E.Z.); and CombGeneTher-HE MSCA Fellowship 101065402 (to E.A.Z.). We thank the Agence Nationale de la Recherche (ANR) IHU FOReSIGHT (ANR-18-IAHU-01) and the I-Demo Regional (BPI, Région Île-de-France) grant GEAR (n^*◦*^ 407307) for funding support.

## 4 Materials and Methods

### 4.1 Datasets

We used two classes of AAV insertion datasets. Per-variant datasets contained sufficiently reliable variant-level viability measurements for supervised model training and evaluation (Table 1). These included two in-house datasets, for AAV2 and AAV5, and one published AAV9 dataset from fit4function [16]. Broad-coverage datasets provided additional sequence coverage or selection outputs but lower per-variant reliability, and were therefore used only for amino-acid enrichment analyses and related aggregate comparisons (Table 2).

### 4.2 In-house experimental procedures

#### AAV production and purification

In-house AAV capsid variant libraries were produced by triple transfection as previously described [42]. Viral particles were purified by iodixanol gradient ultracentrifugation, then concentrated and buffer-exchanged into PBS containing 0.001% Pluronic using Amicon Ultra-50 centrifugal filters (Millipore). All variants packaged the same transgene, carrying a 21-bp barcode at the 3’ end for sequence-level quantification.

#### Post-production inactivation

Purified AAV samples were inactivated with the TruTiter system (Promega) [43]. Briefly, 5 *µ*L of purified AAV was incubated at 37^*◦*^C for 15 minutes with 1 *µ*L viability PCR reagent in a total volume of 100 *µ*L. Then 1 *µ*L neutralization solution was added, followed by 5 minutes at room temperature and 5 minutes at 95^*◦*^C. Inactivated samples were stored at −20^*◦*^C before titration.

#### Sequencing

For the in-house datasets summarized in Table 1, plasmid libraries or inactivated viral DNA were amplified using primers flanking the 7-mer insertion. For AAV5, the primers were AAV5 F (TCGTCGGCAGCGTCAGATGTGTATAA-GAGACAGTCATCACCAGCGAGAGCGAG) and AAV5 R (GTCTCGTGGGCTCGGAGATGTGTATAA-GAGACAGTTTCCTGGAGGTTGTACGTG). For the in-house AAV2 per-variant dataset, the primers were AAV2 F LD (TCGTCGGCAGCGTCAGATGTG-TATAAGAGACAGATCAGGACAACCAATCCCGTG-GCTA) and AAV2 R LD (GTCTCGTGGGCTCGGA-GATGTGTATAAGAGACAGTGTCCTGCCAGAC-CATGCCTG). Libraries were indexed by PCR (8 cycles) using the Nextera XT v2 Index Kit (Illumina), with 10 ng input DNA for plasmid samples or 5 *µ*L of inactivated viral material as input. Sequencing was performed at the BioMics platform (Institut Pasteur, Paris, France) on an Illumina MiSeq100 for the in-house AAV2 per-variant library and on an Illumina NovaSeq X for the AAV5 library, using paired-end 2*×*150 bp reads. PhiX was spiked in at 30% of the total DNA loaded onto the flow cell.

For the in-house datasets summarized in Table 2, the AAV2 plasmid library and viral DNA from viability-selection rounds 1 and 2 were amplified across the CAP insertion site using primers AAV2 F (TATCAGGACAACCAATCCCGTGGCTA) and AAV2 R (CAGGCATGGTCTGGCAGGACA). Amplicons were A-tailed, adapter ligated, and amplified for indexing with the KAPA HyperPrep kit (Kapa Biosystems, Wilmington, MA, USA) according to the manufacturer’s instructions, starting from 10–60 ng DNA. Sequencing was performed at the Genom’IC platform (Institut Cochin, Paris, France) on an Illumina NextSeq 2000 using single-end 1 × 200 bp reads. PhiX was spiked in at 10–25% of the total DNA loaded onto the flow cell.

Across all in-house library preparations, size distributions were assessed with an Agilent Bioanalyzer or TapeStation, DNA concentrations were quantified with the Qubit dsDNA HS assay, and AMPure XP bead purification was performed after each amplification or ligation step. The 7-mer insertion sequences were extracted from NGS reads by locating the constant flanking sequences on both the 5^*′*^ and 3^*′*^ sides of the 21-bp region of interest (using Cutadapt and SeqKit) [44, 45]. Reads were then filtered to retain only sequences with the expected error probability ≤ 0.1 and the correct length (using VSEARCH and `kseq.h`) [46, 47].

### 4.3 Empirical viability and variant filtering

For each variant, sequencing yielded one plasmid-input count and one packaged-viral-pool count. Counts were converted into normalized frequencies after adding a fixed pseudocount (*α* = 1) to each pool independently, and empirical viability was defined as the log_2_ ratio of viral to plasmid abundance:

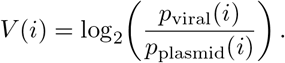

Sampling uncertainty was estimated from the plasmid and viral counts by approximating the variance of a normalized frequency from its count and propagating this uncertainty through the log-ratio. Downstream analyses retained only variants with a viability standard deviation below 0.5, equivalent to Err(*i*) *<* 0.25. Variants failing this filter generally had very low plasmid or viral read counts. Full normalization and uncertainty formulas are provided in the Supplementary Methods.

### 4.4 Viability labels

The empirical viability distributions were bimodal. To define viable and non-viable classes without imposing an arbitrary threshold, we fit a two-component Gaussian mixture model to the filtered viability values using `scikit-learn` (`GaussianMixture`, full covariance, *n*_init_ = 10, regularization 10^−6^). The lower-mean component was assigned as non-viable and the higher-mean component as viable. Model details are provided in the Supplementary Methods.

### 4.5 Biochemical features of inserted peptides

To compare viable and non-viable insertions (Fig. 1f–i), we computed a panel of amino-acid physicochemical properties for each 7-mer. Each insertion was represented by the mean value of each property across its seven residues. The property panel included hydropathy, hydrophilicity, hydrophobicity, bulkiness, side-chain volume, polarity, secondary-structure propensities, flexibility, charge class, aromaticity, molecular weight, and isoelectric point. The full property list and sources are provided in the Supplementary Methods.

### 4.6 Distribution comparisons

To compare viability-score distributions between variant sets (e.g., literature vs. production-selected variants; Fig. 5), we used a weighted Kolmogorov–Smirnov (KS) statistic, defined as the maximum absolute difference between the corresponding weighted empirical cumulative distribution functions. Statistical significance was assessed by bootstrap resampling under the null hypothesis that both samples are drawn from the same underlying viability distribution (Supplementary Methods, Section S2.2).

### 4.7 Baseline sequence and feature models

We used simple baseline models to evaluate how much viability could be explained by coarse biochemical composition, residue position, and low-order sequence interactions.

#### Random forest classifier

A random-forest classifier with 200 trees was trained on averaged biochemical features using the GMM-derived viable and non-viable labels. Model performance was reported as AUC, and feature importance scores were used to summarize which correlated properties contributed most to the classification (Fig. 1j,k).

#### Sequence-based regression models

We trained three regression models directly on 7-mer sequences: a position-independent linear model using amino-acid composition only, a position-dependent linear model using a full 7 by 20 one-hot encoding, and a two-layer multilayer perceptron allowing nonlinear interactions. Models were trained with a variance-weighted mean-squared-error loss, so that high-confidence viability measurements contributed more strongly. Performance was evaluated using Pearson and Spearman correlations between predicted and empirical viability (Fig. 1l). Full model definitions and the loss function are provided in the Supplementary Methods.

### 4.8 PoET zero-shot analyses

PoET is an autoregressive protein language model trained on multiple-sequence alignments and designed to capture evolutionary and structural constraints from protein sequence alone [17]. In all zero-shot analyses, PoET was used without training on AAV insertion viability data.

We used zero-shot PoET scores for two types of analyses. First, we evaluated linker composition by scoring simulated variants in which 7-mer inserts were paired with alternative flanking linkers. Second, we performed capsid-wide in silico insertion scans by introducing random 7-mer insertions at candidate positions and computing both an insertion tolerance score and a structural locality score. For the serotype-transfer analyses, PoET scoring was performed on filtered AAV2, AAV5, and AAV9 datasets after applying the same uncertainty threshold used elsewhere in the manuscript (Err(*i*) *<* 0.25), together with a minimal count filter requiring plasmid and viral counts greater than 1 when these columns were available. This yielded 9,488 AAV2 variants, 49,299 AAV5 variants, and 68,776 AAV9 variants for zero-shot and supervised evaluation. For these scans, PoET scoring is based on sequence log-likelihood; we summarize insertion effects as the change in zero-shot log-likelihood induced by the insertion (relative to the corresponding wild-type background), and report positionwise tolerance values as *z*-scores across VP1 positions to facilitate comparison and visualization. Details of MSA construction, linker simulations, capsid-wide insertions, and locality estimation are provided in the Supplementary Methods.

### 4.9 Supervised prediction with PoET embeddings

When reliable viability measurements were available, we used PoET embeddings as fixed sequence representations for supervised prediction. PoET itself was kept frozen, and only a lightweight prediction head was trained. For each variant, the supervised model used embeddings from the seven inserted residues at the VR-VIII site rather than from a single position. Separate heads were trained for AAV2, AAV5, and AAV9, and cross-serotype transfer was evaluated by applying a head trained on one serotype directly to another. Architecture, optimization, data splits, and evaluation metrics are provided in the Supplementary Methods.

