## Supplementary for "Viability of engineered AAVs via protein language models"

### Supplementary Information for:
*Viability of engineered AAVs via protein language models*

Mélissa Desrosiers[1,†], Tommaso Ocari[1,†], Jeanne Trinquier[1,†], Emília A. Zin[1,†], Timothé Van Meter[1], Maëlle Delmas[1], Müge Tekinsoy[1], Arthur Planul[1], Takahiro Nemoto[2], Deniz Dalkara[1,‡], Ulisse Ferrari[1,‡]
[1] Institut de la Vision, Sorbonne Université, INSERM, CNRS, Paris, France
[2] WPI-PRIMe, Osaka University, Suita, Japan
[†] These authors contributed equally. [‡] These authors contributed equally.
Corresponding authors: Deniz Dalkara, Ulisse Ferrari, and Jeanne Trinquier.
Institut de la Vision, Sorbonne Université, INSERM UMRS 968, Paris, France


#### S1 Supplementary Figures and Tables

##### S1.1 Additional amino acid enrichments

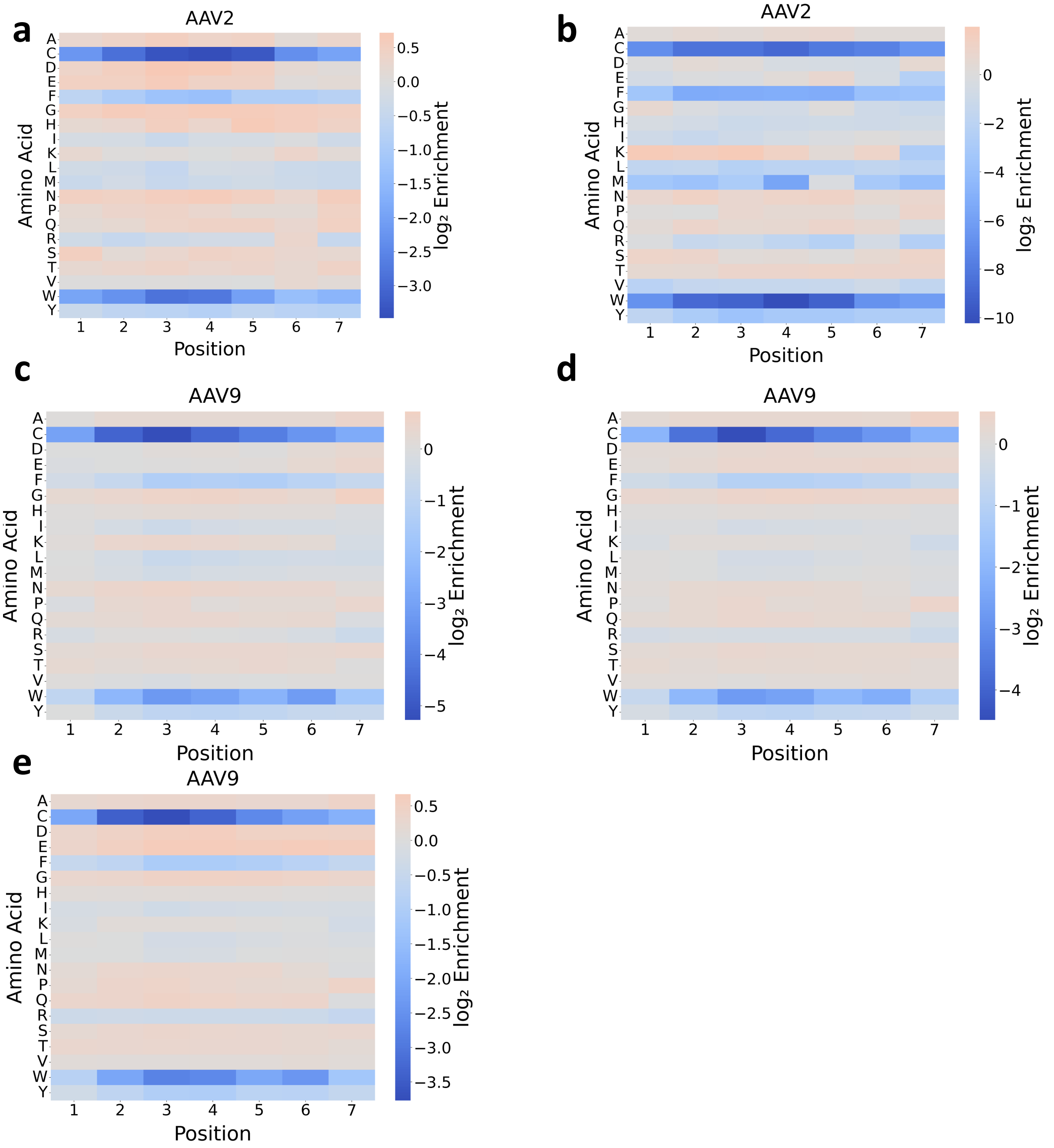


**Supplementary Fig. S1.** Additional amino-acid enrichment profiles across broad-coverage datasets. Panels a to e show position-wise amino-acid enrichments for independent libraries that were used for enrichment analyses only. The datasets correspond to the broad-coverage collection listed in Table 2 of the main text (AAV2 in-house, AAV2 Byrne et al. [1], and three published AAV9 datasets from Stanton et al. [2], Tabebordbar et al. [3], and Ravindra Kumar et al. [4]). Across datasets, the same broad trend is observed: small, flexible, and polar residues are generally enriched, whereas bulky hydrophobic/aromatic residues and cysteine are depleted.

#### S1.2 Linker analysis across serotypes

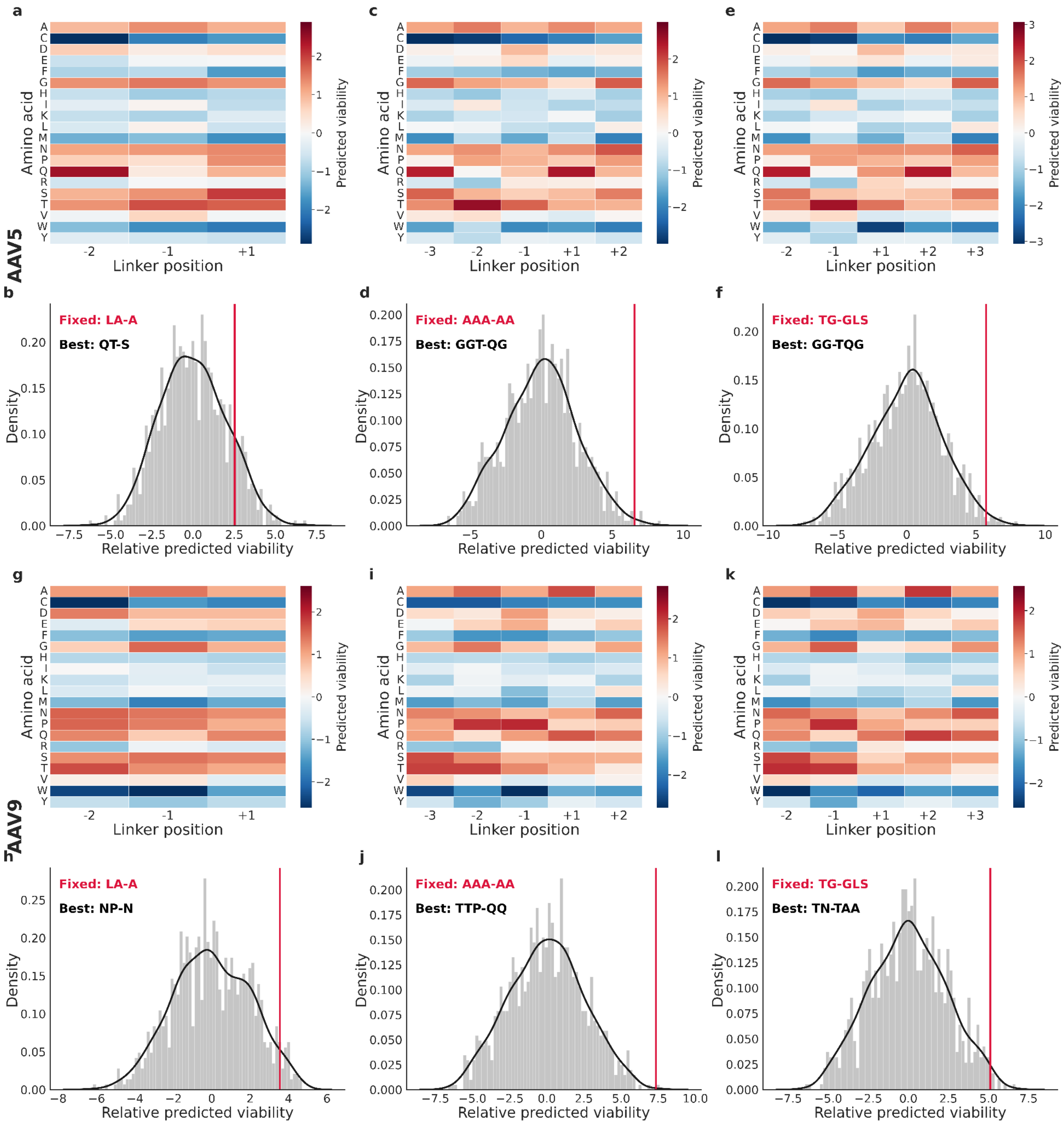


**Supplementary Fig. S2.** Library-level linker analyses stratified by serotype and linker architecture. For AAV5, panels **a,b** correspond to `XX-X`, **c,d** to `XXX-XX`, and **e,f** to `XX-XXX`. For AAV9, the same layout starts at panel **g**: **g,h** for `XX-X`, **i,j** for `XXX-XX`, and **k,l** for `XX-XXX`.

#### S1.3 Insertion tolerance profiles across serotypes

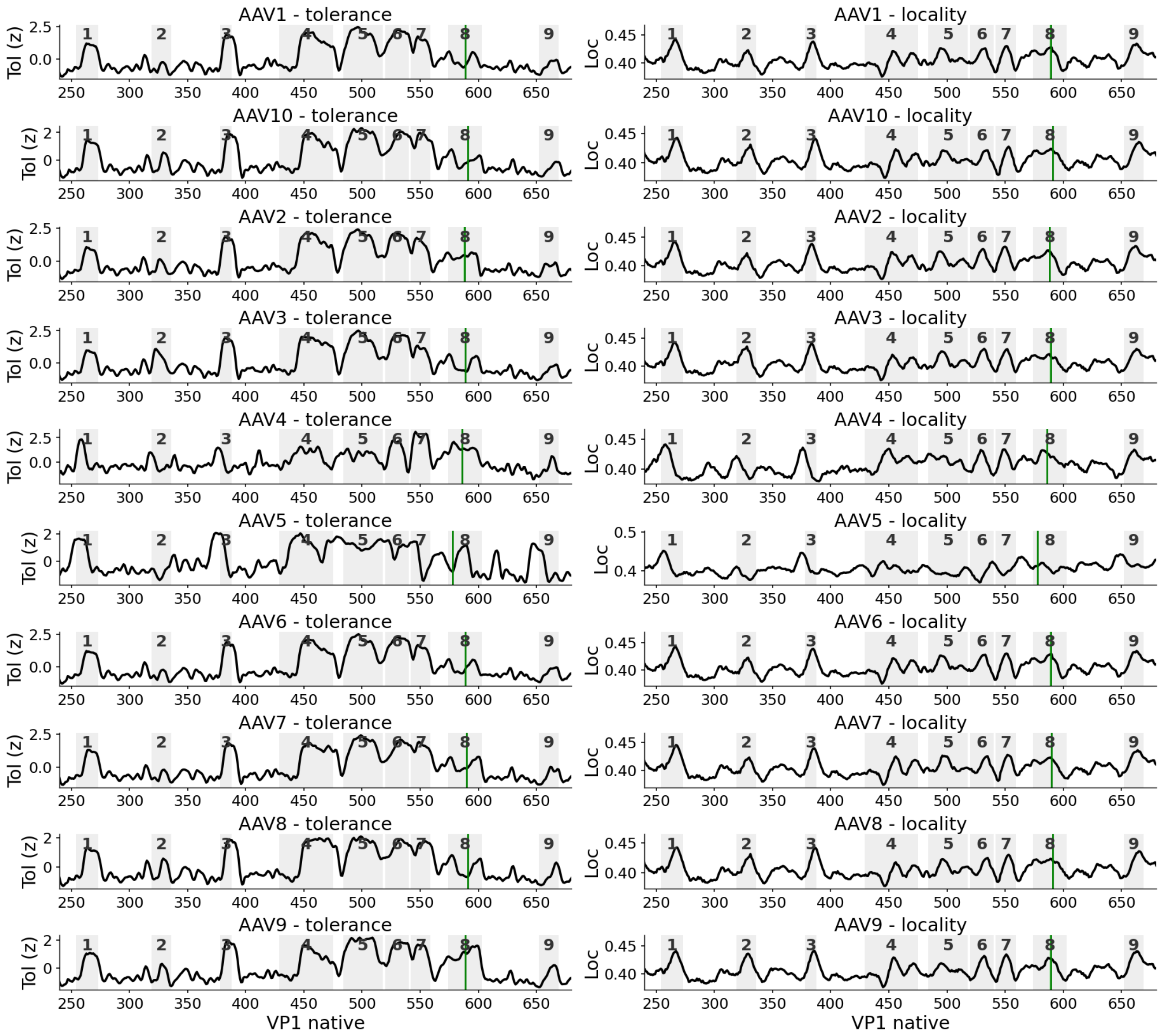


**Supplementary Fig. S3.** Capsid-wide insertion tolerance and locality predictions across serotypes. Panels report per-serotype tolerance/locality profiles used to compute the mean and standard-deviation summaries shown in Fig. 3c,d of the main text. Gray shading marks variable regions and green markers indicate the canonical insertion site.

#### S1.4 Zero-shot and transfer performance summary

**Supplementary Table S1.** Summary of zero-shot, within-serotype, and cross-serotype Spearman correlations. Reported values are the mean, standard deviation, and standard error of the mean across $n = 5$ random seeds.

| **Category** | **Train** | **Test** | $n$ | **Mean** | **SD** | **SEM** |
|---|---|---|---|---|---|---|
| self | AAV2 | AAV2 | 5 | 0.770 | 0.014 | 0.006 |
| self | AAV5 | AAV5 | 5 | 0.412 | 0.013 | 0.006 |
| self | AAV9 | AAV9 | 5 | 0.917 | 0.002 | 0.001 |
| cross | AAV2 | AAV5 | 5 | 0.317 | 0.034 | 0.015 |
| cross | AAV2 | AAV9 | 5 | 0.824 | 0.007 | 0.003 |
| cross | AAV5 | AAV2 | 5 | 0.595 | 0.034 | 0.015 |
| cross | AAV5 | AAV9 | 5 | 0.733 | 0.029 | 0.013 |
| cross | AAV9 | AAV2 | 5 | 0.723 | 0.025 | 0.011 |
| cross | AAV9 | AAV5 | 5 | 0.331 | 0.023 | 0.010 |
| zero-shot | zero-shot | AAV2 | 5 | 0.485 | 0.031 | 0.014 |
| zero-shot | zero-shot | AAV5 | 5 | 0.159 | 0.013 | 0.006 |
| zero-shot | zero-shot | AAV9 | 5 | 0.684 | 0.008 | 0.004 |

### S2 Supplementary Methods

#### S2.1 Empirical viability computation and uncertainty

For each variant $i$, plasmid and viral read counts were first converted into normalized frequencies. A fixed pseudocount $\alpha = 1$ was added to each count to avoid undefined log-ratios for variants with zero reads in one pool:

$$p_{\text{plasmid}}(i) = \frac{c_{\text{plasmid}}(i) + \alpha}{\sum_j \left(c_{\text{plasmid}}(j) + \alpha\right)}, \qquad p_{\text{viral}}(i) = \frac{c_{\text{viral}}(i) + \alpha}{\sum_j \left(c_{\text{viral}}(j) + \alpha\right)}.$$

Empirical viability was defined as

$$V(i) = \log_2\left(\frac{p_{\text{viral}}(i)}{p_{\text{plasmid}}(i)}\right).$$

This score is positive when a variant is enriched in the packaged viral pool relative to the plasmid input, and negative when it is depleted after production. To estimate sampling uncertainty, we approximated the variance of a normalized frequency $p(i)$ as

$$\text{Var}(p(i)) = \frac{1}{c(i) + \alpha}\left[1 - \frac{c(i) + \alpha}{\sum_j \left(c(j) + \alpha\right)}\right],$$

and estimated the variance of viability as

$$\text{Err}(i) = \text{Var}(p_{\text{plasmid}}(i)) + \text{Var}(p_{\text{viral}}(i)).$$

The standard deviation used for filtering in the main text was $\sigma_i = \sqrt{\text{Err}(i)}$.

#### S2.2 Distribution comparisons

Because production-selection datasets are sequencing-based, variants in the selected pool carry different NGS support. We therefore compared predicted-viability distributions using a KS-style distance computed on count-weighted empirical distributions (weights proportional to NGS counts, normalized to sum to 1), while treating literature variants as an unweighted sample.

To obtain a $p$-value, we used bootstrap resampling under the null hypothesis that the two sets are drawn from the same underlying distribution: we pooled the observations, repeatedly resampled two sets of the original sizes with replacement (sampling proportional to weights for the sequencing-based pools), recomputed the KS-style distance for each replicate, and reported the $p$-value as the fraction of replicates with a distance at least as large as the observed one.

#### S2.3 Gaussian mixture modeling of viability

Filtered viability values were separated into viable and non-viable classes using a two-component Gaussian mixture model:

$$p(V) = \pi_1 \mathcal{N}(V \mid \mu_1, \sigma_1^2) + \pi_2 \mathcal{N}(V \mid \mu_2, \sigma_2^2).$$

Parameters were estimated by Expectation–Maximization using `scikit-learn`. Components were ordered by increasing mean, with $\mu_1 < \mu_2$. Variants assigned to the lower-mean component were labeled non-viable, and variants assigned to the higher-mean component were labeled viable.

#### S2.4 Variance-weighted training loss

Regression models were trained with a variance-weighted mean-squared-error loss so that variants with lower measurement uncertainty contributed more strongly:

$$\mathcal{L} = \frac{1}{N}\sum_{i=1}^{N} w(i)\left(\hat{V}(i) - V(i)\right)^2, \qquad w(i) = \frac{1}{\text{Err}(i) + \epsilon},$$

where $\epsilon$ is a small numerical constant.

#### S2.5 Amino-acid property panel

For the amino-acid enrichment analyses and biophysical feature plots, each 7-mer was represented by the average value of each property across its seven residues. The curated property panel comprised Kyte–Doolittle hydropathy, Hopp–Woods hydrophilicity, Eisenberg hydrophobicity, Zimmerman bulkiness, Chothia side-chain volume, Grantham polarity, Chou–Fasman secondary-structure propensities, Bhaskaran–Ponnuswamy flexibility, charge class, aromaticity, molecular weight, and isoelectric point. These properties were implemented as fixed per-amino-acid lookup tables, using standard amino-acid index scales compiled in AAindex [5] when available. Molecular weight and isoelectric point were represented with the approximate residue-level values used in the analysis notebooks.

#### S2.6 Sequence-based baseline models

For the baseline models in Fig. 1l, each inserted peptide was encoded as a $7 \times 20$ one-hot matrix and flattened into a 140-dimensional vector $x(i)$.

**Position-independent linear model (AA).** The one-hot matrix was summed across the seven positions to produce a 20-dimensional amino-acid composition vector $c(i)$. Viability was predicted as

$$\hat{V}(i) = b + \sum_{k=1}^{20} w_k \, c_k(i),$$

where $c_k(i)$ is the number of occurrences of amino acid $k$ in the 7-mer.

**Position-dependent linear model (AA + position).** The position-dependent model used the full one-hot matrix and assigned a coefficient to each position-amino-acid pair:

$$\hat{V}(i) = b + \sum_{p=1}^{7} \sum_{k=1}^{20} w_{p,k} \, x_{p,k}(i).$$

**Multilayer perceptron (AA + position + interactions).** The nonlinear baseline was a two-layer feedforward neural network with ReLU activation:

$$\hat{V}(i) = f_2(\mathrm{ReLU}(f_1(x(i)))) \,,$$

where $f_1$ maps the 140-dimensional input to a 64-dimensional hidden layer and $f_2$ maps the hidden layer to a scalar output.

#### S2.7 PoET zero-shot analyses

All zero-shot PoET analyses were performed on VP1 capsid sequences. Multiple-sequence alignments (MSAs) were constructed with ColabFold using default parameters. In the serotype-transfer notebook, PoET inference used the ColabFold-style alignment file provided at `data/colbfoldmsa/colabfoldoutput.a3m`. To make scores comparable across analyses, PoET inference was run with fixed token limits and a fixed random seed, so that the same MSA subsampling procedure was used across runs. PoET parameters were never updated in these analyses.

**Linker composition.** To estimate general linker preferences, we simulated libraries containing 500 distinct 7-mer insertions. Each insert was paired with 1,000 alternative linker variants of length 3 or 5, depending on the linker layout being evaluated. PoET was used to score the full capsid sequence containing each insert-linker combination. Residue-level linker preferences were obtained by averaging scores across simulated libraries (Fig. 2a–c).

**Library-level linker performance.** For each linker layout, we computed the distribution of mean PoET scores across simulated directed evolution starting libraries (Fig. 2d–f). Common linker motifs (`LA-A`, `AAA-AA`, and `TG-GLS`) were then positioned within these distributions.

**Variant-level linker performance.** For each published insertion variant, we sampled 10,000 alternative linkers of the corresponding length. The linker used in the original construct was ranked against this in silico linker set using its PoET score (Fig. 2h–i).

**Capsid-wide insertion tolerance and locality.** To estimate insertion tolerance across the capsid, we introduced random 7-mer insertions at each candidate position and computed the mean decrease in PoET zero-shot log-likelihood relative to the wild-type sequence (Fig. 3c). Structural locality was estimated from PoET attention maps as the fraction of predicted structural response concentrated within a ±5-residue window around the insertion site (Fig. 3d).

#### S2.8 PoET-supervised viability prediction

For supervised viability prediction, PoET was used as a frozen feature extractor. Forward and backward contextual embeddings at the seven inserted positions were concatenated to form a $7 \times d$ representation. A compact supervised head was trained on top of this representation, consisting of a linear projection, transformer encoder blocks, global average pooling across the seven positions, and a final linear output layer. Only the head parameters were trainable.

The same architecture was used for all serotypes: input dimension $d_{\text{model}} = 2048$, projection to 512 dimensions, transformer encoder with 2 layers and 4 attention heads, feedforward dimension 512, dropout 0.1, and a final scalar output. Optimization used Adam with learning rate $10^{-4}$ and batch size 256 for up to 60 epochs. Training used early stopping on validation loss with patience 5 and minimum improvement $10^{-6}$. The regression loss was the variance-weighted MSE, $(\hat{y} - y)^2/(\text{Err} + 10^{-12})$.

One supervised head was trained per serotype (AAV2, AAV5, and AAV9). Within-serotype performance was evaluated with repeated random splits (train/validation/test = 0.8/0.1/0.1; $n_{\text{seeds}} = 5$). Cross-serotype transfer was evaluated by applying a head trained on one serotype directly to another serotype without retraining. We report the mean and standard deviation of variance-weighted Pearson and Spearman correlations.

#### S2.9 Literature 7-mer insertion variants

**Supplementary Table S2.** Literature-reported AAV variants containing 7-mer insertions analyzed in this study.

| Paper | Serotype | Variant | Linker | Name |
|---|---|---|---|---|
| Jang et al. [6] | AAV2 | TQVGQKT | LA-A | r3.45 |
| Dalkara et al. [7] | AAV2 | LGETTRP | LA-A | 7m8 |
| Dalkara et al. [7] | AAV2 | NETITRP | LA-A | |
| Dalkara et al. [7] | AAV2 | KAGQANN | LA-A | |
| Dalkara et al. [7] | AAV2 | KDPKTTN | LA-A | |
| Ozturk et al. [8] | AAV2 | PDSTTRS | LA-A | |
| Ozturk et al. [8] | AAV2 | PQDTTKK | LA-A | |
| Ozturk et al. [8] | AAV2 | HQDTTKN | LA-A | |
| Ozturk et al. [8] | AAV2 | TTSQNKP | LA-A | |
| Kotterman et al. [9] | AAV2 | ISDQTKH | LA-A | R100 (core 7-mer) |
| Perabo et al. [10] | AAV2 | RGDAVGV | AAA-AA | rAAV-M07A |
| Perabo et al. [10] | AAV2 | RGDTPTS | AAA-AA | rAAV-M07T |
| Perabo et al. [10] | AAV2 | GENQARS | AAA-AA | rAAV-MecA |
| Perabo et al. [10] | AAV2 | RSNAVVP | AAA-AA | rAAV-MecB |
| Chen et al. [11] | AAV2 | WPFYGTP | AAA-AA | |
| Chen et al. [11] | AAV2 | DSPAHPS | AAA-AA | |
| Chen et al. [11] | AAV2 | LPSSLQK | AAA-AA | |
| Chen et al. [11] | AAV2 | GWTLHNK | AAA-AA | |
| Pavlou et al. [12] | AAV2 | GLSPPTR | AAA-AA | AAV2.GL |
| Pavlou et al. [12] | AAV2 | NNPTPSR | AAA-AA | AAV2.NN |
| Pavlou et al. [12] | AAV2 | GAHRSDS | AAA-AA | |
| Pavlou et al. [12] | AAV2 | NSRPAAA | AAA-AA | |
| Pavlou et al. [12] | AAV2 | SSPGLPR | AAA-AA | |
| Rode et al. [13] | AAV2 | THGTPAD | AAA-AA | AAV2-THGTPAD |
| Rode et al. [13] | AAV2 | LPSRPSL | AAA-AA | AAV2-LPSRPSL |
| White et al. [14] | AAV2 | HAIYPRH | | |
| White et al. [14] | AAV2 | THALWHT | | |
| Stiefelhagen et al. [15] | AAV2 | EARVRPP | | |
| Stiefelhagen et al. [15] | AAV2 | NSVSLYT | | |
| Sellner et al. [16] | AAV2 | NDVRSAN | | |

*Continued on next page*

| Paper | Serotype | Variant | Linker | Name |
|---|---|---|---|---|
| Michelfelder et al. [17] | AAV2 | RGDLGLS | | |
| Michelfelder et al. [17] | AAV2 | RGDMSRE | | |
| Michelfelder et al. [17] | AAV2 | DGLGRLV | | |
| Michelfelder et al. [17] | AAV2 | ESGLSQS | | |
| Michelfelder et al. [17] | AAV2 | PRSADLA | | |
| Michelfelder et al. [17] | AAV2 | PRSTSDP | | |
| Michelfelder et al. [17] | AAV2 | ESGLSQS | | |
| Korbelin et al. [18] | AAV2 | NRGTEWD | | AAV-BR1 |
| Korbelin et al. [18] | AAV2 | ADHVQWT | | |
| Korbelin et al. [18] | AAV2 | DDGVSWK | | |
| Korbelin et al. [18] | AAV2 | SDGLTWS | | |
| Korbelin et al. [18] | AAV2 | NNVRTSE | | |
| Korbelin et al. [18] | AAV2 | SDGLAWV | | |
| Korbelin et al. [18] | AAV2 | ESGHGYF | | |
| Korbelin et al. [18] | AAV2 | EYRDSSG | | |
| Korbelin et al. [18] | AAV2 | NDVRAVS | | |
| Deverman et al. [19] | AAV9 | TLAVPFK | | AAV-PHP.B |
| Deverman et al. [19] | AAV9 | YTLSQGW | | AAV-PHP.A |
| Deverman et al. [19] | AAV9 | QAVRTSL | | |
| Ravindra et al. [4] | AAV9 | TLAVPFS | | |
| Ravindra et al. [4] | AAV9 | TALKPFL | | |
| Ravindra et al. [4] | AAV9 | RYQGDSV | | |
| Ravindra et al. [4] | AAV9 | WSTNAGY | | |
| Ravindra et al. [4] | AAV9 | ERVGFAQ | | |
| Stanton et al. [2] | AAV9 | RSVGSVY | | |
| Stanton et al. [2] | AAV9 | KTVGTVY | | |
| Stanton et al. [2] | AAV9 | REQQKLW | | |
| Stanton et al. [2] | AAV9 | PSQGTLR | | |
| Stanton et al. [2] | AAV9 | PTQGTVR | | |
| Stanton et al. [2] | AAV9 | RVDPSGL | | |
| Jang et al. [20] | AAV9 | AGAGPFK | AQ in positions 587 to 588 mutated to DG | |
| Jang et al. [20] | AAV9 | KFPVALT | AQ in positions 587 to 588 mutated to DG | |
| Lin et al. [21] | AAV9 | WPPKTTS | AQ in positions 587 to 588 mutated to DG | |
| Lin et al. [21] | AAV9 | QRPPREP | AQ in positions 587 to 588 mutated to DG | |
| Lin et al. [21] | AAV9 | QRPPRPA | AQ in positions 587 to 588 mutated to DG | |
| Lin et al. [21] | AAV9 | TPPKTTS | AQ in positions 587 to 588 mutated to DG | |
| Lin et al. [21] | AAV9 | EPPKTTS | AQ in positions 587 to 588 mutated to DG | |
